# RNA3DB: A structurally-dissimilar dataset split for training and benchmarking deep learning models for RNA structure prediction

**DOI:** 10.1101/2024.01.30.578025

**Authors:** Marcell Szikszai, Marcin Magnus, Siddhant Sanghi, Sachin Kadyan, Nazim Bouatta, Elena Rivas

## Abstract

With advances in protein structure prediction thanks to deep learning models like AlphaFold, RNA structure prediction has recently received increased attention from deep learning researchers. RNAs introduce substantial challenges due to the sparser availability and lower structural diversity of the experimentally resolved RNA structures in comparison to protein structures. These challenges are often poorly addressed by the existing literature, many of which report inflated performance due to using training and testing sets with significant structural overlap. Further, the most recent Critical Assessment of Structure Prediction (CASP15) has shown that deep learning models for RNA structure are currently outperformed by traditional methods.

In this paper we present RNA3DB, a dataset of structured RNAs, derived from the Protein Data Bank (PDB), that is designed for training and benchmarking deep learning models. The RNA3DB method arranges the RNA 3D chains into distinct groups (Components) that are non-redundant both with regard to sequence as well as structure, providing a robust way of dividing training, validation, and testing sets. Any split of these structurally-dissimilar Components are guaranteed to produce test and validations sets that are distinct by sequence and structure from those in the training set. We provide the RNA3DB dataset, a particular train/test split of the RNA3DB Components (in an approximate 70/30 ratio) that will be updated periodically. We also provide the RNA3DB methodology along with the source-code, with the goal of creating a reproducible and customizable tool for producing structurally-dissimilar dataset splits for structural RNAs.

**Graphical Abstract:** 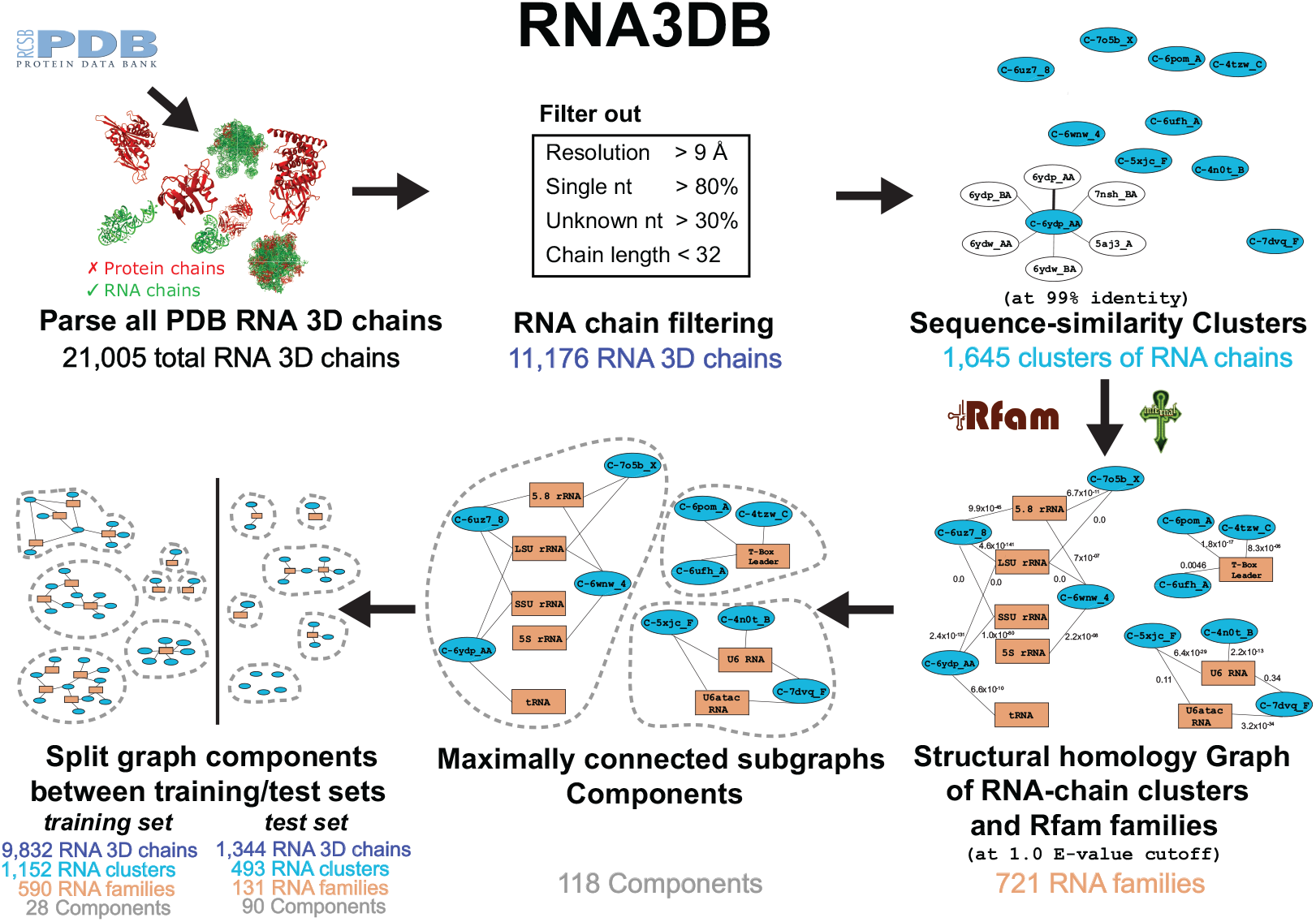

**Highlights:**

- While there is a recent surge in applying deep learning to RNA structure prediction, domain experts have raised concerns about generalization and current trends in benchmarking.
- Many of the concerns primarily relate to how novel RNA families–i.e. families unseen in the training set–are benchmarked, and whether the models are effective at handling such cases. Performance on bench-marks reflective of real-world applications, such as CASP15 and RNA-Puzzles, is poor for RNA deep learning models.
- We present a dataset–RNA3DB–that is designed for training and bench-marking deep learning models for RNA structure prediction. RNA3DB provides coverage of all RNA chains found in the Protein Data Bank (PDB).
- RNA3DB is clustered into groups that are both sequentially and structurally non-redundant, providing a robust way of creating training, validation, and testing sets for deep learning models. Along with the dataset, we also provide a transparent methodology as well as the source-code, making our tool both reproducible and customizable.

## 1. Introduction

Since DeepMind introduced AlphaFold [1] at CASP13 in 2018, there has been growing interest in applying deep learning to problems in structural biology [2]. When AlphaFold2 improved on its predecessor’s already impressive results in 2020, many jumped to claim that protein structure prediction was “solved” [3].

Naturally it didn’t take long for lessons from AlphaFold to start being adapted for RNA [4, 5, 6, 7, 8, 9, 10, 11, 12], with some papers immediately reporting impressive results both for RNA secondary and tertiary structure prediction. Certainly at first glance, RNA structure appears to be in some ways analogous to protein structure. In both cases, the one-dimensional polymer sequences fold into three-dimensional conformations which are strongly tied to the molecule’s function, and strongly dependent on the molecule’s sequence.

Despite this, most RNA biologists do not consider deep learning methods to be state-of-the-art for RNA structure prediction [13] for secondary or tertiary structures. Publications began pointing out issues with generalization to unseen sequences in deep learning methods for secondary structure prediction in 2022 [14, 15, 16, 17], although this overfitting behavior was already well-known for probabilistic and Markov random field models trained on structural RNA data since 2012 [18]. With the increased interest in RNA, partly attributed to AlphaFold’s success, and partly due to the rise of RNA-based therapeutics [19], CASP15 joined RNA-Puzzles [20] and added 12 RNA-only targets to the competition [21, 22], where all deep learning methods performed comparatively poorly to traditional tools [23]. Since CASP15, several other publications have attempted to apply deep learning to the RNA problem including DeepMind [24], many of whom report impressive results [23], but often without addressing the concerns regarding generalization.

There is a widespread belief in deep learning that the quantity and quality of datasets is one of the most influential aspects towards a model’s performance. The number of available proteins is significantly higher than RNAs in the Protein Data Bank (PDB). Figure 1 shows a comparison between the number of protein and RNA experimentally determined structures deposited in PDB (named chains). After filtering (see Section 3.2), PDB contains nearly 70 times more available protein chains than RNA chains.

**Figure 1.**
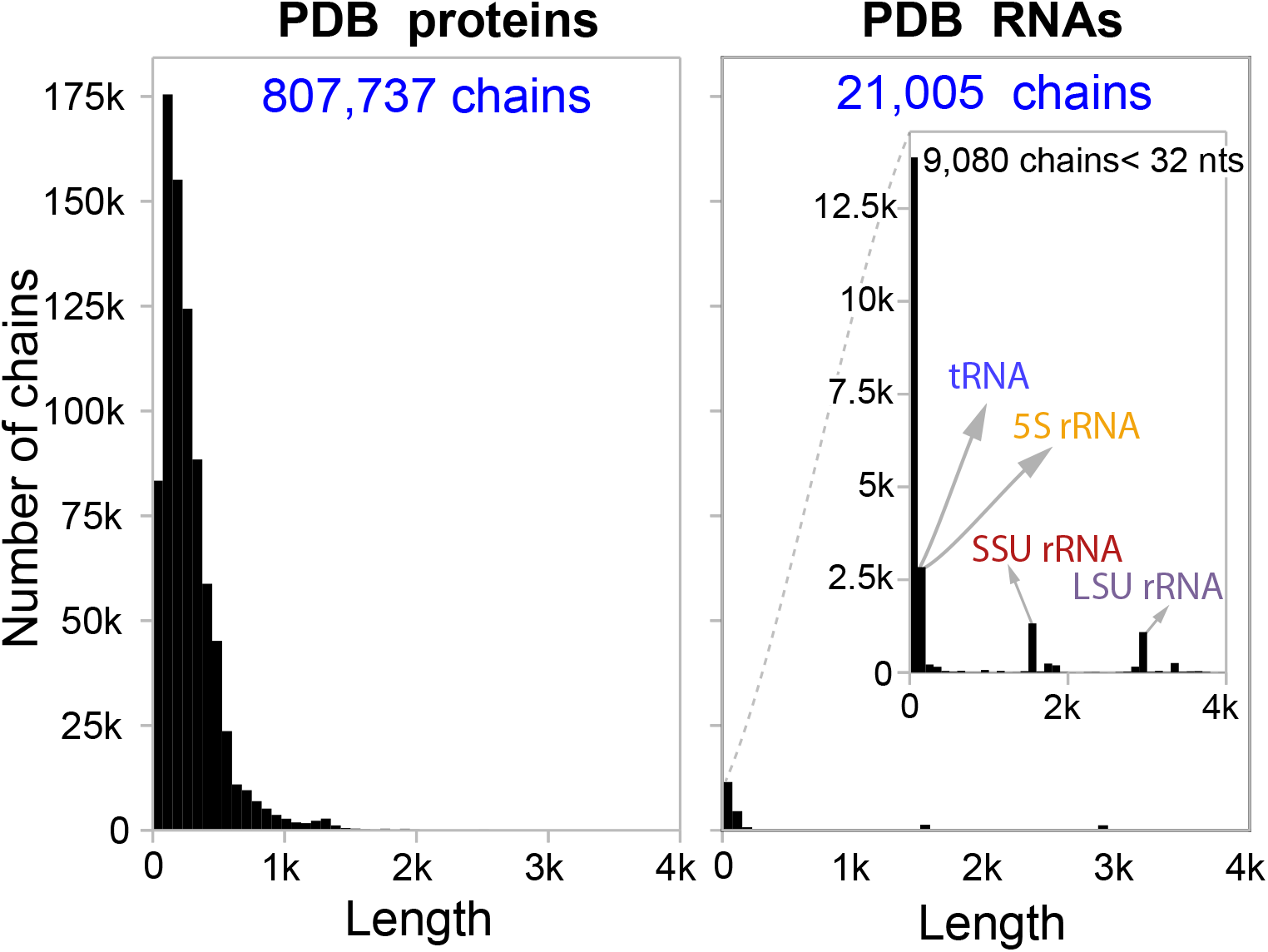
Comparison of length distributions of chains in the PDB for proteins and RNAs at the same y-axis scale. The inset plot shows a zoomed in version of the RNA histogram for visibility.

With this in mind, it is perhaps no surprise that deep learning for RNA is less successful than for proteins. While other problems are potentially solvable engineering challenges, the quantity and diversity of available data cannot be easily increased. Fortunately, the number of novel structures uploaded to the PDB each years appears to be increasing (Figure 2), albeit still very far away from proteins. It is important to note that the PDB, while comprehensive and consistently used by the structural community, is a repository, and not a dedicated tool to provide deep learning models with a curated dataset, leading to several challenges. For instance, many of the RNAs included in the PDB are short fragments of longer RNAs, and many of the entries contain the same or effectively the same RNA sequences repeatedly.

**Figure 2.**
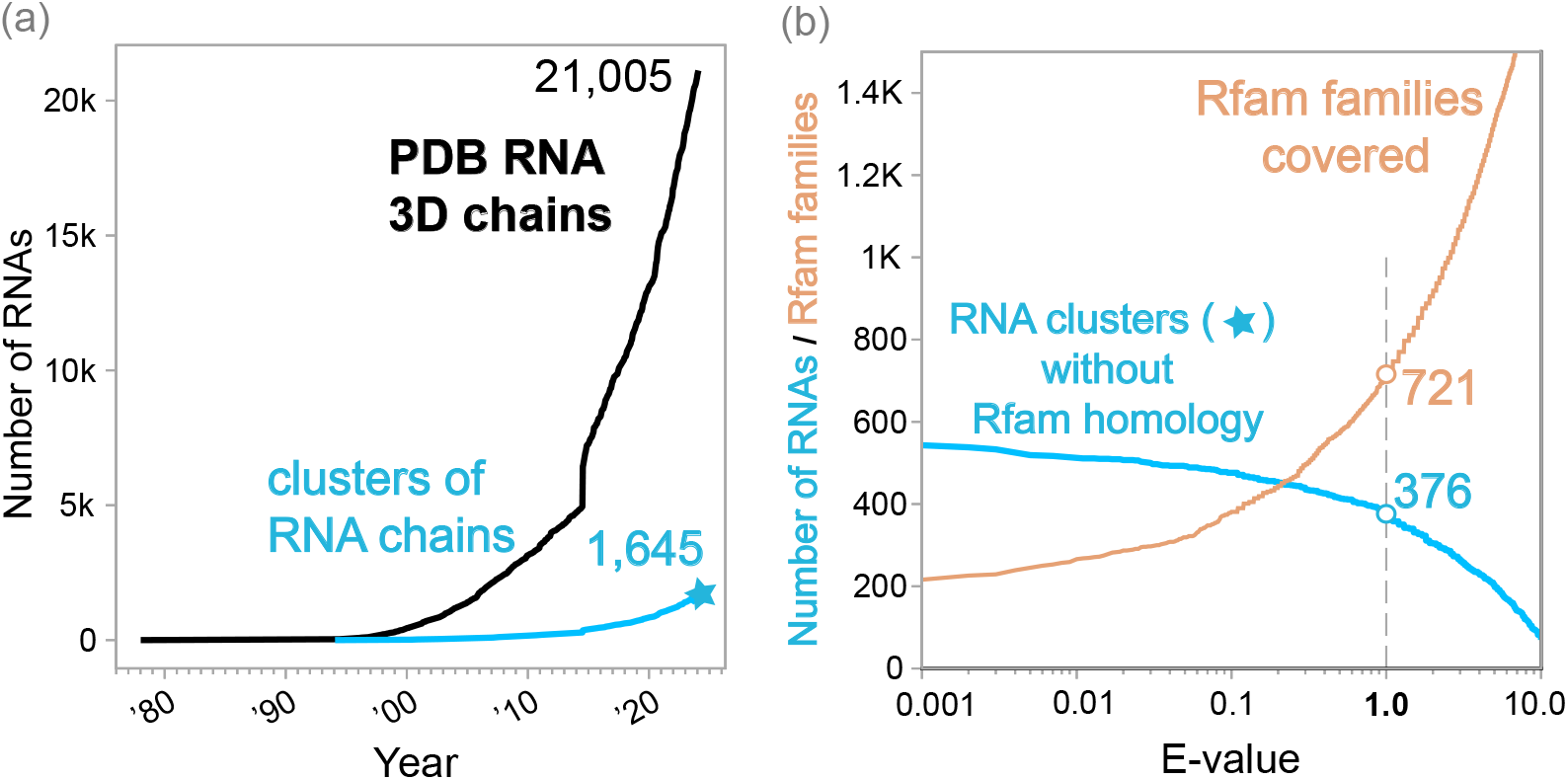
Panel **(a)** shows: in black the cumulative number of PDB chains that contain at least one RNA residue based on the `_chem_comp.type` data item by year; and in blue the cumulative number of distinct RNA sequences (sequence identity threshold of 99%) represented in the PDB RNA 3D chains. Panel **(b)** shows in blue the number of distinct RNA sequences in the PDB as of 2024-01-10 (blue star in (a)) without significant homology to any Rfam structural RNA family at different E-value thresholds (as calculated by the Infernal method `cmscan`). Panel **(b)** shows in orange the diversity of Rfam RNA structures covered by the RNA chains at different E-value cutoffs.

RNA3DB is a dataset based on all PDB RNA 3D structures developed for addressing the aforementioned concerns, especially those regarding generalization, particularly for training and benchmarking deep learning models. The methodology behind RNA3DB clusters the RNA 3D structures into distinct groups that are non-redundant both with regard to sequence as well as structure, providing a robust way of dividing training, validation, and testing sets.

## 2. Comparison to other methods

There are many existing methods that process the PDB and extract RNA chains in order to produce databases that capture the diversity of existing RNA 3D structures, while also removing redundancies present in PDB [25, 26, 27, 28, 29, 30, 31, 32]. These efforts have been useful for many years to the RNA community.

The advent of methods for RNA secondary structure prediction trained on data (either by refining thermodynamic parameters [33], or by training exclusively from known RNA structures [34, 35, 18]) has brought the need to construct datasets that go beyond just characterizing the existing RNA 3D structures to the attention of the community. In addition, there is a need for datasets that are able to provide rigorous separation of the data into structurally dissimilar sets. To our knowledge, this separation by dissimilar structures found in the PDB RNA chains is not provided by existing tools. The structural separation is needed in order to distinguish whether the methods are able to generalize to new structures or not; otherwise data leakage results in overfitting [18]. Recent studies of deep learning methods have also shown the need to separate structures between training and testing sets [14, 15, 16, 17, 23]. The goal of RNA3DB is to provide such structurally dissimilar grouping and splitting of the existing RNA 3D structures present in PDB.

Amongst the existing methods, the RNA 3D Hub [27] introduces a non-redundant dataset of RNA 3D structures, each of which represents an “equivalence classe”. These equivalence classes are meant to avoid the redundancy inherent in PDB due to multiple 3D structures of the same molecule from the same organism, but the system is not designed to assure that structures in different classes do not share any structural homology. For instance, the human and fly 28S+5.8S rRNAs are two different equivalence classes in RNA 3D Hub of the same RNA structures. Other datasets such as RNAnet [30] and RNAsolo [31] are based on this RNA 3D Hub structurally-redundant set of representative RNA structures.

Within structural RNAs, structural similarity can be often identified amongst sequences that have low sequence similarity. However, databases such as RNA-NRD [32] considers that chains with high structural similarity but low sequence identity (*<*80%) are not redundant. As a result, of the 3,175 RNA-NRD clusters, for instance, 589 correspond to tRNA or tRNA elements, as tRNAs through covariation conserve the clover-leaf structure at the expense of low sequence similarity. Many other RNA-NRD clusters also share structural similarity with each other. On the other hand, RNA3DB uses Infernal to assess the structural similarity for every chain against all the Rfam models and makes sure that chains with any possibility of being structurally similar, regardless of sequence identity are grouped together in any train/test dataset split.

Some methods provide Rfam family information [27, 30, 32], but do not use it as a part of their identification of “non-redundant” RNAs. This is sufficient for many applications, but not for training and benchmarking deep learning models. Even for databases that provide Rfam family information, creating structurally dissimilar groups is not trivial, since as previously pointed out by Szikszai *et al*. [14], naively separating by Rfam families is insufficient. There are many Rfam families that share homology. Rfam groups families that show some homology to each other into Clans. However, Rfam Clans or even higher level classifications based on homology, such as the “Superfamilies” built on top of Clans in the method RNArchitecture [29], are insufficient for splitting RNA chains into structurally dissimilar groups. It is not uncommon to find entries in PDB that combine non-homologous structural RNAs in one chain, for instance rRNA and tRNAs (see `PDB:6YDP_AA` [36, 37]). In fact, in our homology search of unique sequences (99% similarity), we find that 10 chains overlap multiple Clans (Supplementary Table S1), and 44 chains overlap multiple families that do not belong to a given Clan (Supplementary Table S2), at an E-value of 10^*−*3^. At an E-value of 1.0, 52 RNA chains overlap multiple Clans, and 460 overlap multiple families without a common Clan.

RNA3DB has been developed to address these issues. Structural families with Infernal [38] homology hits to the same RNA chains are grouped together into a Component during the graph building step (Section 3.4). Each Component is a collection of Rfam families and PDB RNA chains that share some level of structural homology (determined by a E-value cutoff set to a lenient value to avoid false negatives). Different Components are guaranteed to be non-redundant with respect to both sequence and importantly also structure. The generation of these Components (Section 3.4.2), which can then be safely assigned to training, testing, or even validation sets without risking data leakage (Section 3.5) are the novel contributions that differentiate RNA3DB from any existing method.

Notice that while RNA 3D Hub and derived databases select one arbitrary RNA chain per equivalence class, RNA3DB saves all the sequence-clustered RNA chains. While the chain is the same, it is possible that the presence of different interacting partners in the actual crystal structures may result in different structural conformations. RNA3DB preserves the full extent of the structural diversity present in PDB.

## 3. Materials and Methods

Our methodology for building RNA3DB can be broken down into five main steps: Parsing, Filtering, Clustering, identification of structurally-dissimilar Components, and Splitting. During the parsing step, all PDB chains are processed to identify potential RNAs. Filtering then removes sequences that are unsuitable for deep learning for various reasons, such as length or resolution. Next, Clustering assigns PDB chains into two hierarchies of groups based on sequence similarity and then structural similarity. After Clustering, a graph of structural homology between all RNA chains to all RNA structural families allows us to identify structurally dissimilar groups, named graph Components. Finally, Splitting of the graph Components assigns PDB chains into training and testing sets that are non-redundant both in terms of sequence and structure.

### 3.1 Parsing

Our method starts with careful parsing of all entries in the PDB to identify any potential RNA structures. This is done by downloading a copy of all PDB entries in PDBx/mmCIF format. To avoid reliance on author labeling, we scan all chains for the `chem_comp` data category, which is found in practically all PDBx/mmCIF files [39]. During our first pass, we accept any chains with at least one residue containing “RNA” in its `_chem_comp.type` data item. Any non-RNA residues, such as amino acids, are treated as “unknown” residues at this stage.

An important parsing issue is that of RNA residue modifications such as pseudouridylations. The number of modified residues in an RNA chain can be large. As an example, for the tRNA structure `PDB:1EHZ` [40, 41] 2 out of the 72 residues are pseudouridines, and 10 are other modified residues. A naive parsing method usually reports these modifications as “unknown” residues.

The RNA3DB method systematically converts all modified nucleic acids to their closest one-letter symbols. We extract all three-letter symbol conversions of nucleic acids (and proteins for possible future use) from the Chemical Component Dictionary, which is an “external reference file describing all residue and small molecule components found in PDB entries” [42]. In addition to naively converting three-letter codes, we also recursively parse parent components from `_chem_comp.mon_nstd_parent_comp_id` to maximize the number of extracted modifications. Any three-letter codes that cannot be converted (including stray amino acids) are parsed as “unknown” residues. Our method is comprehensive method, and is able to identify up to 582 different nucleic acid modifications.

### 3.2 Filtering

Next, the Filtering step aims to remove chains that are not informative for training deep learning models. By default, RNA3DB considers four filtering categories: sequence length, structural resolution, fraction of individual nucleotides in the sequence, and fraction of “unknown” residues.

Chains shorter than 32 residues are removed during this step. In many cases, there is not much information about the structure in only a few nucleotides, and these sequences can be largely ignored. However, it should be noted that this is not always true. There are also some cases where some of these shorter chains are potentially informative fragments of longer RNAs (for instance, chain `PDB: 354D_A`^1^ [43, 44] is a crystal structure exclusively of the 12 nucleotide long Loop E of 5S rRNA). Since these short motifs are difficult to classify into families–as due to their short length they are hard to distinguish from random sequences–we opt to remove these from the dataset as it may lead to data leakage. We experimented with methods to attempt to both identify and keep these fragments, but were unable to systematically avoid scenarios where known fragments of longer chains overlapped between training and testing sets.

Chains with structural resolution higher than 9Å are removed, as they are considered to have too low confidence to be useful to determine atom positions. This is the same threshold that AlphaFold2 uses [1]. By default RNA3DB also excludes any structures resolved with nuclear magnetic resonance spectroscopy (NMR), as NMR does not provide well-defined resolution values. However this only excludes a relatively small number (177) of chains that would otherwise not be removed. An optional flag exists within RNA3DB’s parser that interprets the resolution of NMR structures as 0.0Å, which allows these structures to be included if desired.

The sequence nucleotide composition is also considered to avoid repetitive sequences with low information content. By default, we remove any sequence where a single nucleotide makes up more than 80% of residues, like AlphaFold2 [1].

Finally, we also remove any sequence where more than 30% of the residues are “unknown”. This removes chains that do not provide sufficient information in their sequences, but also acts as a filter that removes any special cases where a non-RNA polymer is parsed as one because the sequence may contain “RNA” in its `_chem_comp.type` (see Section 3.1).

### 3.3 Clustering

Clustering is divided into two distinct steps: sequence-based Clustering, and structure-based Clustering.

#### 3.3.1 Sequence-based Clustering

First, MMseqs2 [45] is used to cluster all 3D RNA chains by 99% sequence identity. Many of the RNA chains in the PDB have identical or near-identical sequences, and the purpose of this step is to group them together in clusters. All RNA chains in one cluster have almost identical RNA sequences, but each chain is associated to a different experimentally determined structure. A given cluster may contain RNA chains identical to a region of a larger chain. RNA3DB selects the longest RNA chain as representative of the cluster, which is named as `“cluster <chain_name>“`. We observe (Figure 2a) that the number of RNA-chain sequence-based clusters is about a tenth that of the number of actual RNA chains.

#### 3.3.2 Structure-based Clustering

Second, Infernal [38] is used to run an RNA homology search against all Rfam families [46]. We find it convenient to have this information for all existing sequences in the PDB, however, this step can be restricted to only the unique sequences after sequence-based Clustering for the purpose of building RNA3DB. We use a two-pass approach to maximize the number of chains for which we get at least one hit, regardless of significance. The first pass uses default Infernal parameters. The second pass re-runs the search on sequences without any hits, except with all filters turned off. The purpose of this second pass is to increase sensitivity, but it is generally very slow.

### 3.4 Structurally dissimilar groups of RNA 3D structures

RNA3DB uses the sequence-clustered RNA chains and the extensive structural homology searches to create structurally dissimilar groups of RNA 3D structures. These steps create a “hierarchy” in RNA3DB, starting with RNA chains, to sequence-based “Clusters”, followed by structure-based independent “Components”. Each Cluster guarantees that its sequences are not identical (or near-identical) to any other Clusters’, while each Component guarantees that it shares no structural homology to the same RNA families as any other Component.

#### 3.4.1 A Graph of RNA chains and structural families

A graph is constructed as follows: let each cluster of RNA chains be a node in the graph, and all Rfam families also be nodes in the graph. Edges are undirected and unweighted between chain and family nodes and are present when some E-value threshold is met. By default we use a generous E-value threshold of 1.00, since we want to eliminate the possibility of missing any potential homology, with false positives being an acceptable trade-off.

#### 3.4.2 Maximally connected subgraphs (Graph Components)

Finally, the RNA3DB method performs a depth-first search on the graph described above to identify all maximally connected subgraphs, named Components. This way, we guarantee that all Components share no homology to the same families. These Components are then ranked by size (i.e. number of unique chain clusters in the Component) with the exception of `Component #0`, which includes all chain-nodes without edges to any RNA family nodes.

We can motivate building these structurally dissimilar Components via some clear examples. Take a 55S mammalian mitochondrial ribosome like `PDB:6YDP_AA` [36, 37]. Infernal finds significant hits (at an E-value threshold of 10^*−*9^) to both LSU and SSU rRNA families, as well as several other hits to tRNAs within the sequence. While rRNA and tRNA families have no homology, this chain must be used in the same training/testing set with other individual tRNA and rRNA chains to avoid inadvertently leaking structural information from the training set into the testing set.

It may be surprising to find out that for 376 chains in the PDB, i.e. those in `Component #0`, we are unable to find homology to any Rfam family at an E-value threshold of 1.0. Many of these chains are synthetically designed structures, messenger RNA fragments crystallized as part of translational complexes, or in some cases structural fragments that are too short, even above 32 residues, to classify at the desired threshold.

### 3.5 Splitting

This is the final step, which assigns the graph Components into training and testing sets. Since any two Components of the graph (Section 3.4) are completely non-redundant, the Components can safely be placed arbitrarily into any set without data leakage.

RNA3DB provides an algorithm for dividing the graph Components into training/test sets. The algorithm simply assigns Components with the largest number of unique sequences into the training set until a specified training set split percentage is met. By default, we recommend a split of 70-30, or in other words, include the largest Components in the training set (with the exception of `Component #0`) until at least 70% of the data is in the training set.

We recommend using `Component #0` for testing rather than training to minimize the chance of data leakage, as well as following structure prediction benchmarking best practices, particularly with regards to reporting results per family instead of an overall average [47]. Alternatively, RNA3DB gives the option to ignore this `Component #0` all together.

It should be noted that this Splitting step can also be done manually with relative ease. Using default parameters, RNA3DB finds 118 graph Components, which is a manageable set for manual inspection.

The RNA3DB database derived from PDB is been updated consistently every three months. In addition, the RNA3DB code is open source and available to produce custom Components and splits for any custom collection of RNA 3D structures.

## 4. Results

Among the most important observations from our dataset is that approximately 1 in 10 RNA PDB chains are either redundant in sequence or too short to be usable by deep learning methods (Figure 2a). Despite this, it is clear that the number of novel RNAs uploaded has increased over recent years.

The RNA3DB parser finds 21,005 RNAs in the PDB as of 2024-01-10. The length filter removes 9,080 chains, while the resolution filter removes 1,540 chains. We find 1,294 sequences dominated by one nucleotide, and 177 that have too many unknown residues to keep. Note that a single chain may be rejected by more than one filter. After filtering 11,176 RNA chains remain, and 9,829 chains are rejected.

Next, the RNA3DB method produces 1,645 sequence-similarity clusters (at 99% identity) of RNA chains. The largest cluster with 629 RNA chains is the *Thermus thermophilus HB8* 70S ribosome. The median number of chains per cluster is 2.0.

Then RNA3DB proceeds to make a graph which adds 721 Rfam family nodes (at an E-value threshold of 1.0) to the 1,645 RNA-chain cluster nodes (Figure 2b). The RNA3DB resulting graph has a total of 3,994 edges. The tRNA (`RF00005`) family has the largest number of edges (307), and the median number of edges per RNA family node is 2.0. The cluster with the largest number of edges (43) is `cluster 6ydp_AA (the 55S mammalian mitochondrial ribosome)`, and the median number of edges per cluster node is 2.0.

Finally, the RNA3DB method produces 118 non-redundant Components. The largest Component, `Component #1`, includes 119 RNA families and 935 RNA-chain clusters. More than half of the Components include one single RNA family and one single RNA cluster of chains.

The `Component #0` set comprises all RNA chains without hit to any Rfam family. At an E-value threshold of 1.0, `Component #0` contains 376 RNA clusters and 979 actual RNA chains (Figure 2b), and it includes synthetic RNAs as well as small messenger RNA sequences crystallized as part of larger complexes.

The RNA3DB dataset provides a training/test dataset Split described in Table 1. The training/testing mmCIF files for the chains in both sets (after converting modified residues) can be downloaded directly from https://github.com/marcellszi/rna3db/releases/latest/.

**Table 1:** Hierarchical table of a training/testing Split. A partial representation of all tiers of hierarchy (Components, sequences, chains and RNA families) for both training and testing sets is shown. RNA3DB uses by default an Infernal E-value cutoff of 1.0 to generate the graph.

| RNA3DB | Graph<br>Components | RNA<br>sequences | RNA<br>chains | RNA families<br>E-val < 10<sup>-3</sup> | RNA families<br>E-val < 1.0 | Description |
| --- | --- | --- | --- | --- | --- | --- |
| **Training** | 28 | 1,152 | 9,832 | 169 | 590 |  |
|  | Component #1 | 935 | 9,037 | 119 | 508 | rRNA (LSU,SSU,5.8,5S),<br>tRNA, tmRNA, U1, etc. |
|  | Component #2 | 24 | 24 | 24 | 24 | Purine/2dG-II riboswitches |
|  | Component #3 | 22 | 82 | 2 | 5 | tracrRNA, CRISPR-DR22 |
|  | Component #4 | 18 | 69 | 2 | 3 | Group I intron |
|  | : |  |  |  |  | : |
|  | Component #28 | 3 | 11 | 1 | 1 | Bacteriophage pRNA |
| **Testing** | 90 | 493 | 1,344 | 47 | 131 |  |
|  | Component #0 | 376 | 979 | 0 | 0 | synthetic RNAs, mRNAs, etc. |
|  | Component #29 | 3 | 12 | 1 | 1 | THF riboswitch |
|  | Component #30 | 3 | 12 | 1 | 1 | ZMP-ZTP riboswitch |
|  | : |  |  |  |  | : |
|  | Component #116 | 1 | 2 | 0 | 1 | L25-Gammaproteobacteria<br>ribosomal protein leader |
|  | Component #117 | 1 | 1 | 0 | 1 | mir-3135 microRNA precursor |
| **Total** | 118 | 1,645 | 11,176 | 216 | 721 |  |

## 5. Implementation

The database and code for RNA3DB can be found at: https://github.com/marcellszi/rna3db along with documentation on the Github repository’s Wiki at https://github.com/marcellszi/rna3db/wiki.

For the RNA3DB dataset provided with this manuscript, we used Rfam version 14.10, Infernal version 1.1.4, and we included all RNA chains from PDB as of 2024-01-10. The Filtering, Clustering, Component and Splitting process, with the exception of the homology search, can be run on a 10 core Apple M2 Pro in under 2 minutes. The homology search took 110 hours on a single Intel Xeon Platinum 8358 processor with 32 cores.

The RNA3DB database will be updated every three months. RNA3DB updates affected solely by a PDB update require only to add edges for the new RNA chains. In this case, an Infernal homology search must be done on the new chains only, which in most cases can be done in minutes. Less frequent updates tied to an Rfam version release require the recalculation of all the edges for the whole graph by running the Infernal homology search for all chains against the new Rfam models. This step currently takes approximately 110 hours.

The RNA3DB method can be customized to build specialized train/test datasets. For instance, Filtering parameters (such as the minimal length) can be modified. Different train/test independent Splits can be created depending on the specified parameters, which are documented in the Github Wiki.

## 6. Discussion

The development of a standardized dataset of RNA structures [48] specifically targeting deep learning, is extremely valuable. Here, we introduce RNA3DB a method to obtain comprehensive information about structured RNAs in the PDB in a format that organizes the RNA chains by their structural homology. RNA3DB builds datasets of maximally connected RNA chains with information on the actual structures represented in each group. RNA3DB exhaustively parses all modified residues present in the experimental RNA chains, and the method is customizable in a number of ways such as minimal length or sequence complexity. RNA3DB makes the reduced amount of structural RNA data present in the PDB apparent when compared to that of proteins, and the limited set of distinct 3D RNA structures that it represents.

The sparsity of data alone could be responsible for the poor performance of deep learning methods dependent on millions of parameters to predict RNA structure, as it was seen at CASP15 [21, 22] for 3D structure prediction, as well as for 2D structure prediction [14, 15, 16, 17]. Other factors possibly handicapping the prediction of RNA structure are the more complex RNA backbone geometry that involves more atoms and degrees of freedom, as well as global nature of RNA secondary structure. The global nature of the secondary structure, unlike with proteins, cannot be inferred locally from contiguous residues, and it substantially informs the 3D structure.

Nevertheless, the sparsity of data deserves prioritized consideration. Would a method trained with very limited data be able to generalize to describe not seen before RNA structures? The realm of image data analysis seems to suggest that deep learning methods are able to generalize even when the amount of training data can just be memorized by the method [49, 50]. The RNA3DB method and its final outcome the RNA3DB dataset is a comprehensive classification of structurally dissimilar RNA experimentally-determined structures. We hope that this tool will provide the RNA structure modeling community an effective tool to investigate this question under different settings with rigor.

## 7. Acknowledgments

We thank Rfam and Blake Sweeney for the helpful early discussions on building RNA3DB. This work was supported by U.S. National Institutes of Health grants (1R01GM144423, and 1R21GM148902 to Elena Rivas).

## Footnotes

1 Chain naming is done via author labeling.

