## Supplementary material for "RNA3DB: A structurally-dissimilar dataset split for training and benchmarking deep learning models for RNA structure prediction": plot_figure2.ipynb

```
# %% [code]
from matplotlib.backends.backend_pgf import FigureCanvasPgf
from matplotlib.dates import YearLocator, DateFormatter
from matplotlib.ticker import FuncFormatter
from datetime import datetime

import matplotlib.transforms as mtransforms
import matplotlib.pyplot as plt

import numpy as np
import matplotlib


matplotlib.backend_bases.register_backend('pdf', FigureCanvasPgf)
matplotlib.rcParams.update({
    "pgf.texsystem": "pdflatex",
    'font.family': 'serif',
    'font.size' : 11,
    'text.usetex': True,
    'pgf.rcfonts': False,
})

LINE_INCHES = 5.39749
ASPECT_RATIO = 4/3
PLOT_WIDTH = LINE_INCHES
PLOT_HEIGHT = PLOT_WIDTH / ASPECT_RATIO

# %% [code]
# load part (a) data
x_parse, y_parse = [], []
with open('figure_2a_data_parse.txt') as f:
    for line in f:
        date, num = line.split()
        x_parse.append(datetime.fromisoformat(date))
        y_parse.append(int(num))

x_cluster, y_cluster = [], []
with open('figure_2a_data_cluster.txt') as f:
    for line in f:
        date, num = line.split()
        x_cluster.append(datetime.fromisoformat(date))
        y_cluster.append(int(num))

# %% [code]
# load part (b) data
x_eval, num_chains, num_fams = [], [], []
with open('figure_2b_data.txt') as f:
    next(f) # skip header
    for line in f:
        line = line.split(',')
        x_eval.append(float(line[0]))
        num_chains.append(int(line[1]))
        num_fams.append(int(line[2]))

# %% [code]
def human_format(num):
    num = float('{:.3g}'.format(num))
    magnitude = 0
    while abs(num) >= 1000:
        magnitude += 1
        num /= 1000.0
    return '{}{}'.format('{:f}'.format(num).rstrip('0').rstrip('.'), ['', 'K', 'M', 'B', 'T'][magnitude])

# %% [code]
fig, ax = plt.subplots(1, 2)
fig.set_size_inches(w=PLOT_WIDTH*1.5, h=PLOT_HEIGHT*1.0)
trans = mtransforms.ScaledTranslation(-20/72, 7/72, fig.dpi_scale_trans)

# PLOT (a)
# actually plot
ax[0].plot_date(x_parse, np.cumsum(y_parse), '-', label='Total', color='k')
ax[0].plot_date(x_cluster, np.cumsum(y_cluster), '-', label='Unique', color='deepskyblue')
ax[0].plot(x_cluster[-1], np.cumsum(y_cluster)[-1], '*', color='deepskyblue')
# setup axes
ax[0].xaxis.set_major_locator(YearLocator(10))
ax[0].xaxis.set_minor_locator(YearLocator(1))
ax[0].xaxis.set_major_formatter(DateFormatter('\'%y'))
ax[0].get_yaxis().set_major_formatter(FuncFormatter(lambda x, p: human_format(x)))
ax[0].set_xlabel('Year')
ax[0].set_ylabel('Number of RNAs')
# various annotations
ax[0].text(0.0, 1.0, '(a)', transform=ax[0].transAxes + trans, va='bottom')
ax[0].annotate('PDB RNA\n3D chains', fontsize=9, xy=(datetime.fromisoformat('2000-01-01'), 17000), color='k', ha='center')
ax[0].annotate(f'{np.cumsum(y_parse)[-1]:,}', fontsize=9, xy=(datetime.fromisoformat('2012-01-01'), np.cumsum(y_parse)[-1]-700), color='k')
ax[0].annotate('RNA-chain\nsequences', fontsize=9, xy=(datetime.fromisoformat('2000-01-01'), 6000), color='deepskyblue', ha='center')
ax[0].annotate(f'{np.cumsum(y_cluster)[-1]:,}', xy=(datetime.fromisoformat('2015-01-01'), np.cumsum(y_cluster)[-1]+1000), color='deepskyblue')
# date format stuff
ax[0].autoscale_view()
fig.autofmt_xdate()

# PLOT (b)
# actually plot
ax[1].plot(x_eval, num_chains, color='deepskyblue')
ax[1].vlines(1.0, 0, 1000, 'k', '--', zorder=0)
# setup axes
ax[1].get_yaxis().set_major_formatter(FuncFormatter(lambda x, p: human_format(x)))
ax[1].set_xlabel('E-value')
ax[1].set_xscale('log')
ax[1].set_ylabel('Number of RNAs')
ax[1].set_xlim((1e-3, 10.0))
ax[1].set_ylim((0, 1500))
# various annotations 
ax[1].text(0.0, 1.0, '(b)', transform=ax[1].transAxes + trans, va='bottom')
ax[1].plot(0.41, 468, '*', color='deepskyblue', zorder=2)
ax[1].annotate('RNA-chain seqs ( )\nwithout\nRfam homology', fontsize=9, xy=(0.03, 390), color='deepskyblue', ha='center', zorder=1)
ax[1].annotate('Rfam families\ncovered', fontsize=9, xy=(0.02, 250), color='darkgoldenrod', ha='center')
ax[1].plot(1, 376, '*', color='deepskyblue')
ax[1].annotate('376', fontsize=9, xy=(2.1, 480), color='deepskyblue', ha='center')
# colour y-axis
ax[1].tick_params(axis='y', colors='deepskyblue')
ax[1].yaxis.label.set_color('deepskyblue')
# twin axis
ax_in = ax[1].twinx()
# actually plot
ax_in.plot(x_eval, num_fams, color='darkgoldenrod')
ax_in.plot(1, 721, '*', color='darkgoldenrod')
# setup axes
ax_in.set_ylabel('Number of Rfam families')
ax_in.set_ylim((0, 1500))
ax_in.tick_params(axis='y', colors='darkgoldenrod')
ax_in.get_yaxis().set_major_formatter(FuncFormatter(lambda x, p: human_format(x)))
# colour axis
ax_in.spines['right'].set_color('darkgoldenrod')
ax_in.spines['left'].set_color('deepskyblue')
ax_in.yaxis.label.set_color('darkgoldenrod')
# annotate
ax_in.annotate('721', fontsize=9, xy=(0.33, 1000), color='darkgoldenrod', ha='center')

fig.tight_layout()
fig.savefig('Figure_2.pdf')

# %% [code]


# %% [code]
```
