## Supplementary material for "RNA3DB: A structurally-dissimilar dataset split for training and benchmarking deep learning models for RNA structure prediction": components_parse.pl

```
#!/usr/bin/perl -w
# components_parse.pl

use strict;
use Class::Struct;

# find directory where the script is installed

use vars qw ($opt_v );  # required if strict used
use Getopt::Std;
use Cwd qw(cwd);

getopts ('v');

# Print a helpful message if the user provides no input file.
if (!@ARGV) {
        print "usage:  components_parse.pl [options] <componentsfile>  \n\n";
        print "options:\n";
 	exit;
}

my $componentsfile = shift;

my $n_comp = 0;
my @name_comp;
my @clust_per_comp;
my @chain_per_comp;
my @fam_per_comp;
my @famatone_per_comp;
parse_components($componentsfile, \$n_comp, \@name_comp, \@clust_per_comp, \@chain_per_comp, \@fam_per_comp, \@famatone_per_comp);

print "n_components $n_comp\n";


# edges per clusters
my $total_clust    = 0;
my $total_chain    = 0;
my $total_fam      = 0;
my $total_famatone = 0;
my %clust_hash;
my %chain_hash;
my %fam_hash;
my %famatone_hash;
for ( my $c = 0; $c < $n_comp; $c ++) {
    push @{$clust_hash{$clust_per_comp[$c]}},       $name_comp[$c];
    push @{$chain_hash{$chain_per_comp[$c]}},       $name_comp[$c];
    push @{$fam_hash{$fam_per_comp[$c]}},           $name_comp[$c];
    push @{$famatone_hash{$famatone_per_comp[$c]}}, $name_comp[$c];
    $total_clust    += $clust_per_comp[$c];
    $total_chain    += $chain_per_comp[$c];
    $total_fam      += $fam_per_comp[$c];
    $total_famatone += $famatone_per_comp[$c];
}

print "\nclusters     $total_clust\n";
my $ncomp = 0;
for my $clust_per_comp ( sort {$b <=> $a} keys %clust_hash ) {
    print "clusters per comp $clust_per_comp\n";
    
    foreach my $name_comp ( @{$clust_hash{$clust_per_comp}} ) {
	$ncomp ++;
        print "        $ncomp $name_comp\n";
	if ($ncomp == int($n_comp/2)|| $ncomp == int($n_comp/2)+1) {
	    print "Median clusters $ncomp/$n_comp at $clust_per_comp\n";
	}
    }
}

print "\nchains       $total_chain\n";
$ncomp = 0;
for my $chain_per_comp ( sort {$b <=> $a} keys %chain_hash ) {
    print "chains per comp $chain_per_comp\n";
    
    foreach my $name_comp ( @{$chain_hash{$chain_per_comp}} ) {
	$ncomp ++;
        print "        $ncomp $name_comp\n";
	if ($ncomp == int($n_comp/2)|| $ncomp == int($n_comp/2)+1) {
	    print "Median chains $ncomp/$n_comp at $chain_per_comp\n";
	}
    }
}

print "\nfams         $total_fam\n";
$ncomp = 0;
for my $fam_per_comp ( sort {$b <=> $a} keys %fam_hash ) {
    print "fams per comp $fam_per_comp\n";
    
    foreach my $name_comp ( @{$fam_hash{$fam_per_comp}} ) {
	$ncomp ++;
        print "        $ncomp $name_comp\n";
	if ($ncomp == int($n_comp/2)|| $ncomp == int($n_comp/2)+1) {
	    print "Median fams $ncomp/$n_comp at $fam_per_comp\n";
	}
    }
}

print "\nfams at one  $total_famatone\n";
$ncomp = 0;
for my $famatone_per_comp ( sort {$b <=> $a} keys %famatone_hash ) {
    print "fams at one per comp $famatone_per_comp\n";
    
    foreach my $name_comp ( @{$famatone_hash{$famatone_per_comp}} ) {
	$ncomp ++;
        print "        $ncomp $name_comp\n";
	if ($ncomp == int($n_comp/2)|| $ncomp == int($n_comp/2)+1) {
	    print "Median fams at one $ncomp/$n_comp at $famatone_per_comp\n";
	}
    }
}


sub parse_components {
    my ($components_file, $ret_n_comp, $name_comp_ref, $clust_per_comp_ref, $chain_per_comp_ref, $fam_per_comp_ref, $famatone_per_comp_ref) = @_;

    my $n_comp = 0;
    
    open (F, "$components_file") || die;
    while(<F>) {
	if (/^\s*(component_\d+)\s+(\S+)\s+(\S+)\s+(\S+)\s+(\S+)\s*$/) {
	    my $component = $1;
	    my $nclust    = $2;
	    my $nchain    = $3;
	    my $nfam      = $4;
	    my $nfamatone = $5;

	    $name_comp_ref->[$n_comp]         = $component;
	    $clust_per_comp_ref->[$n_comp]    = $nclust;
	    $chain_per_comp_ref->[$n_comp]    = $nchain;
	    $fam_per_comp_ref->[$n_comp]      = $nfam;
	    $famatone_per_comp_ref->[$n_comp] = $nfamatone;
	    $n_comp ++;
	}
    }
    close(F);

    $$ret_n_comp = $n_comp;

}
```
