## Supplementary material for "RNA3DB: A structurally-dissimilar dataset split for training and benchmarking deep learning models for RNA structure prediction": find_edges.pl

```
#!/usr/bin/perl -w
# find_edges.pl

use strict;
use Class::Struct;

# find directory where the script is installed

use vars qw ($opt_v );  # required if strict used
use Getopt::Std;
use Cwd qw(cwd);

getopts ('v');

# Print a helpful message if the user provides no input file.
if (!@ARGV) {
        print "usage:  find_edges.pl [options] <graphfile>  \n\n";
        print "options:\n";
 	exit;
}

my $splitfile = shift;
my $graphfile = shift;

my $n_chain = 0;
my $n_clust = 0;
my @num_chain_per_clust;
my @name_chain_per_clust;
my @name_clust;
parse_split($splitfile, \$n_chain, \$n_clust, \@num_chain_per_clust, \@name_chain_per_clust, \@name_clust);
print "From split: n_cluster $n_clust n_chain $n_chain\n";

my @edges_per_clust;
for (my $c = 0; $c < $n_clust; $c ++) {
    $edges_per_clust[$c] = 0;
}

my $n_fam = 0;
my @name_fam;
my @edges_per_fam;
parse_graph_for_edges($graphfile, $n_clust, \@name_clust, \@edges_per_clust, \$n_fam, \@name_fam, \@edges_per_fam);


# edges per clusters
my $edges_total_clust = 0;
my %clust_hash;
for ( my $c = 0; $c < $n_clust; $c ++) {
    push @{$clust_hash{$edges_per_clust[$c]}}, $name_clust[$c];
    $edges_total_clust += $edges_per_clust[$c];
}

my $ncl = 0;
print "\nEDGES per cluster, n_clust $n_clust\n";
for my $e_per_cl ( sort {$b <=> $a} keys %clust_hash ) {
    print "edges per cluster $e_per_cl\n";
    
    foreach my $name_c ( @{$clust_hash{$e_per_cl}} ) {
	$ncl ++;
        print "        $ncl $name_c\n";
	if ($ncl == int($n_clust/2)|| $ncl == int($n_clust/2)+1) {
	    print "Median $ncl/$n_clust at $e_per_cl\n";
	}
    }
}

# edges per family with hits
my $edges_total_fam   = 0;
my %fam_hash;
for ( my $f = 0; $f < $n_fam; $f ++) {
    push @{$fam_hash{$edges_per_fam[$f]}}, $name_fam[$f];
    $edges_total_fam += $edges_per_fam[$f];
}

if ($edges_total_clust != $edges_total_fam) {
    print "n edges does not match. by clust $edges_total_clust by fam $edges_total_fam\n";
    exit();
}
print "total # edges $edges_total_clust\n";

my $nfam = 0;
print "\nEDGES per fam with hit, n_fam $n_fam\n";
for my $e_per_fam ( sort {$b <=> $a} keys %fam_hash ) {
    print "edges per fam $e_per_fam\n";
    
    foreach my $name_f ( @{$fam_hash{$e_per_fam}} ) {
	$nfam ++;
        print "        $nfam $name_f\n";
	if ($nfam == int($n_fam/2)|| $nfam == int($n_fam/2)+1) {
	    print "Median $nfam/$n_fam at $e_per_fam\n";
	}
    }
    
}
    

    
sub parse_graph_for_edges {
    my ($graph_file, $n_clust, $name_clust_ref, $edges_per_clust_ref, $ret_n_fam, $name_fam_ref, $edges_per_fam_ref) = @_;

    my $n_fam = 0;
    
    open (F, "$graph_file") || die;
    while(<F>) {
	if (/^component_(\d+)\s+(\S+)\s(\S+)\s+(\S+)\s+(\S+)\s*$/) {
	    my $component = $1;
	    my $thisclust = $2;
	    my $thisfam   = $3;

	    my $nc = 0;
	    for (my $n = 0; $n < $n_clust; $n ++) {
		if ($thisclust =~ /^$name_clust_ref->[$n]$/) {
		    $edges_per_clust_ref->[$n] += 1;
		    $nc ++;
		}
	    }
	    if ($nc != 1) 
	    {
		print "found cluster $thisclust $nc times, should be one\n"; exit();
	    }

	    my $f  = 0;
	    my $nf = 0;
	    for ($f = 0; $f < $n_fam; $f ++) {
		if ($thisfam =~ /^$name_fam_ref->[$f]$/) {
		    $edges_per_fam_ref->[$f] += 1;
		    $nf ++;
		}
	    }
	    if ($f == $n_fam && $nf == 0) 
	    {
		$name_fam_ref->[$f]      = $thisfam;
		$edges_per_fam_ref->[$f] = 1;
		$n_fam += 1;
		$nf    += 1;
	    }
	    if ($nf != 1) 
	    {
		print "found fam $thisfam $nf times, should be one\n"; exit();
	    }
	
	}
    }
    close(F);

    $$ret_n_fam = $n_fam;
}


sub parse_split {
    my ($split_file, $ret_n_chain, $ret_n_clust, $num_chain_per_clust_ref, $name_chain_per_clust_ref, $name_clust_ref) = @_;

    my $n_chain = 0;
    my $n_clust = 0;

    my $prv_cluster = "";
    
    open (F, "$split_file") || die;
    while(<F>) {
	if (/^(\S+)\s+(\S+)\s+(\S+)\s+(\S+)\s+(\S+)\s+(\S+)\s+(\S+)\s*$/) {
	    my $whichtest = $1;
	    my $component = $2;
	    my $cluster   = $3;
	    my $chain     = $4;
	    my $res       = $5;
	    my $date      = $6;
	    my $sequence  = $7;

	    if ($cluster =~ /^$prv_cluster$/) {
		$num_chain_per_clust_ref->[$n_clust-1]  += 1;
		$name_chain_per_clust_ref->[$n_clust-1] .= $chain."-";
	    }
	    else {
		$n_clust ++;
		$num_chain_per_clust_ref->[$n_clust-1]  = 1;
		$name_chain_per_clust_ref->[$n_clust-1] = $chain."=";
		$name_clust[$n_clust-1] = $cluster;
		$prv_cluster = $cluster;
	    }
	    $n_chain ++;
	}
    }
    close(F);
    
    $$ret_n_chain = $n_chain;
    $$ret_n_clust = $n_clust;
    
}
```
