## Supplementary material for "RNA3DB: A structurally-dissimilar dataset split for training and benchmarking deep learning models for RNA structure prediction": graph_parse.pl

```
#!/usr/bin/perl -w
#pdb_parse.pl

use strict;
use Class::Struct;

# find directory where the script is installed

use vars qw ($opt_v );  # required if strict used
use Getopt::Std;
use Cwd qw(cwd);

getopts ('v');

# Print a helpful message if the user provides no input file.
if (!@ARGV) {
        print "usage:  graph_parse.pl [options] <graphfile>  \n\n";
        print "options:\n";
 	exit;
}

my $graphfile = shift;

my $n_cluster = 0;
my @cluster;
my @which_fam;
my @which_comp;

parse_graph($graphfile, \$n_cluster, \@cluster, \@which_fam, \@which_comp);

print "n_cluster $n_cluster\n";
for (my $n = 0; $n < $n_cluster; $n ++) {
    print "$which_comp[$n] ($n) $cluster[$n]\n$which_fam[$n]\n";
    #print "$which_comp[$n] ($n) $cluster[$n] $which_fam[$n]\n";
}

sub parse_graph {
    my ($graph_file, $ret_n_cluster, $cluster_ref, $which_fam_ref, $which_comp_ref) = @_;

    my $n_cluster = 0;
    
    open (F, "$graph_file") || die;
    while(<F>) {
	if (/^component_(\d+)\s+(\S+)\s(\S+)\s+(\S+)\s+(\S+)\s*$/) {
	    my $component = $1;
	    my $thiscluster = $2;
	    my $Rfam      = $3."=".$4."=".$5;

	    my $n = 0;
	    for ($n = 0; $n < $n_cluster; $n ++) {
		if ($thiscluster =~ /^$cluster_ref->[$n]$/) {
		    $which_fam_ref->[$n] .= $Rfam."\n";
		    #$which_fam_ref->[$n] .= $Rfam."  ";
		    last;
		}
	    }
	    if ($n == $n_cluster) 
	    {
		$cluster_ref->[$n]    = $thiscluster;
		$which_fam_ref->[$n]  = $Rfam."\n";
		#$which_fam_ref->[$n]  = $Rfam." ";
		$which_comp_ref->[$n] = $component;
		$n_cluster += 1;
	    }
		
	}
    }
    close(F);

    $$ret_n_cluster = $n_cluster;

}
```
