## Supplementary material for "RNA3DB: A structurally-dissimilar dataset split for training and benchmarking deep learning models for RNA structure prediction": split_parse.pl

```
#!/usr/bin/perl -w
# splt_parse.pl

use strict;
use Class::Struct;

# find directory where the script is installed

use vars qw ($opt_v );  # required if strict used
use Getopt::Std;
use Cwd qw(cwd);

getopts ('v');

# Print a helpful message if the user provides no input file.
if (!@ARGV) {
        print "usage:  split_parse.pl [options] <splitfile>  \n\n";
        print "options:\n";
 	exit;
}

my $splitfile = shift;

my $n_chain = 0;
my $n_clust = 0;

my $N     = 1000;
my $k     = 1;
my $shift = 0;

my @chain_per_clust;
my @name_clust;

parse_split($splitfile, \$n_chain, \$n_clust, \@chain_per_clust, \@name_clust);

my $nchain = 0;
my $nchain_max = -1;
for (my $c = 0; $c < $n_clust; $c ++) {
    my $cur_chains = $chain_per_clust[$c];
    $nchain += $cur_chains;
    #printf "clust %d $name_clust[$c] nchain $cur_chains\n", $c+1;
}
print "split $n_chain/$nchain $n_clust\n";
if ($n_chain != $nchain) { print "chains is $nchain should be $n_chain\n"; exit; }

my %hash;
for ( my $c = 0; $c < $n_clust; $c ++) {
    push @{$hash{$chain_per_clust[$c]}}, $name_clust[$c];
}

my $ncl = 0;
for my $ch_per_cl ( sort {$b <=> $a} keys %hash ) {
    print "nchain per cluster $ch_per_cl\n";
    
    foreach my $name_c ( @{$hash{$ch_per_cl}} ) {
	$ncl ++;
        print "        $ncl $name_c\n";
	if ($ncl == int($n_clust/2)|| $ncl == int($n_clust/2)+1) {
	    print "Median $ncl/$n_clust at $ch_per_cl\n";
	}
    }
}
    

sub parse_split {
    my ($split_file, $ret_n_chain, $ret_n_clust, $chain_per_clust_ref, $name_clust_ref) = @_;

    my $n_chain = 0;
    my $n_clust = 0;

    my $prv_cluster = "";
    
    open (F, "$split_file") || die;
    while(<F>) {
	if (/^(\S+)\s+(\S+)\s+(\S+)\s+(\S+)\s+(\S+)\s+(\S+)\s+(\S+)\s*$/) {
	    my $whichtest = $1;
	    my $component = $2;
	    my $cluster   = $3;
	    my $chain     = $4;
	    my $res       = $5;
	    my $date      = $6;
	    my $sequence  = $7;

	    if ($cluster =~ /^$prv_cluster$/) {
		$chain_per_clust_ref->[$n_clust-1] ++;
	    }
	    else {
		$n_clust ++;
		$chain_per_clust_ref->[$n_clust-1] = 1;
		$name_clust[$n_clust-1] = $cluster;
		$prv_cluster = $cluster;
	    }
	    $n_chain ++;
	}
    }
    close(F);
    
    $$ret_n_chain = $n_chain;
    $$ret_n_clust = $n_clust;
    
}
```
