## Supplementary figures and images for "RNA3DB: A structurally-dissimilar dataset split for training and benchmarking deep learning models for RNA structure prediction"

### Figure_2.pdf

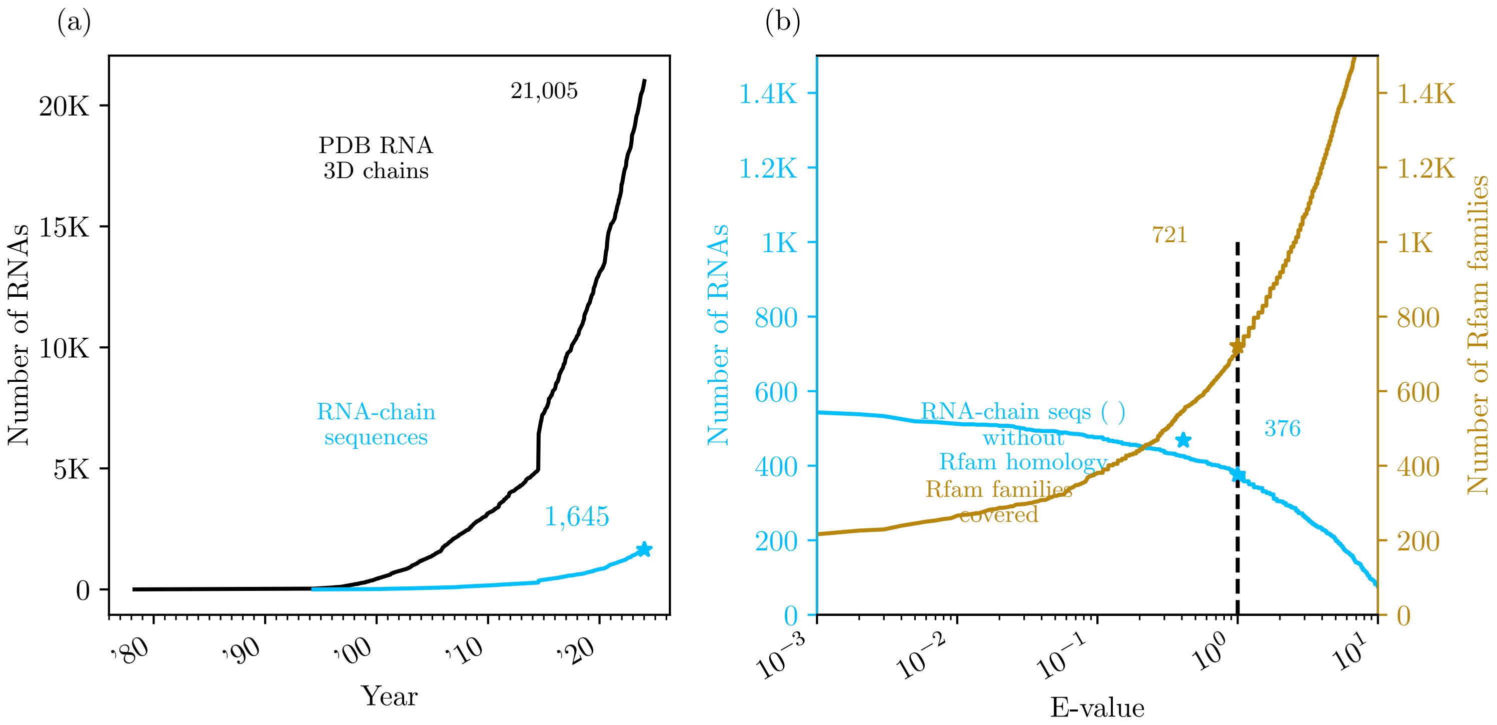
