## supplemental tables for "RNA3DB: A structurally-dissimilar dataset split for training and benchmarking deep learning models for RNA structure prediction"

#### Contents

**1 Table of PDB RNA chains with homology to multiple Rfam Clans 2**

**2 Table of PDB RNA chains with homology to multiple Rfam families that do not share a common Rfam Clan 3**

*Preprint submitted to J. Mol. Biol.*

### 1. Table of PDB RNA chains with homology to multiple Rfam Clans

**Table S1:** PDB RNA chain cluster representatives (at 99% sequence similarity) with homology to multiple Rfam Clans. We provide all Infernal `cmscan` hits at an E-value threshold of $10^{-3}$.

| RNA chain | Start | End | Rfam accession | Name | Clan accesion | Clan name | E-value |
|---|---|---|---|---|---|---|---|
| 7am2_1 | 1,904 | 2,470 | RF02546 | LSU_trypano_mito | CL00112 | LSU | 1.4e-138 |
| 7am2_1 | 2,491 | 3,117 | RF02545 | SSU_trypano_mito | CL00111 | SSU | 4e-129 |
| 6hrm_1 | 1,507 | 4,403 | RF02541 | LSU_rRNA_bacteria | CL00112 | LSU | 0.0 |
| 6hrm_1 | 1,506 | 4,411 | RF02540 | LSU_rRNA_archaea | CL00112 | LSU | 0.0 |
| 6hrm_1 | 1 | 1,528 | RF00177 | SSU_rRNA_bacteria | CL00111 | SSU | 0.0 |
| 6hrm_1 | 1,666 | 4,414 | RF02543 | LSU_rRNA_eukarya | CL00112 | LSU | 0.0 |
| 6hrm_1 | 6 | 1,460 | RF01959 | SSU_rRNA_archaea | CL00111 | SSU | 3.5e-274 |
| 6hrm_1 | 6 | 1,457 | RF02542 | SSU_rRNA_microsporidia | CL00111 | SSU | 8.3e-199 |
| 6hrm_1 | 6 | 1,522 | RF01960 | SSU_rRNA_eukarya | CL00111 | SSU | 8.4e-181 |
| 6hrm_1 | 1,519 | 1,672 | RF00002 | 5_8S_rRNA | CL00112 | LSU | 2.5e-07 |
| 6hrm_1 | 4,372 | 4,445 | RF00177 | SSU_rRNA_bacteria | CL00111 | SSU | 0.00072 |
| 7aor_2 | 2,020 | 2,579 | RF02546 | LSU_trypano_mito | CL00112 | LSU | 9.7e-138 |
| 7aor_2 | 2,589 | 3,210 | RF02545 | SSU_trypano_mito | CL00111 | SSU | 4.5e-120 |
| 6wnw_4 | 1 | 3,031 | RF02541 | LSU_rRNA_bacteria | CL00112 | LSU | 0.0 |
| 6wnw_4 | 1 | 3,031 | RF02540 | LSU_rRNA_archaea | CL00112 | LSU | 0.0 |
| 6wnw_4 | 160 | 3,021 | RF02543 | LSU_rRNA_eukarya | CL00112 | LSU | 0.0 |
| 6wnw_4 | 1,069 | 1,185 | RF00001 | 5S_rRNA | CL00113 | 5S_rRNA | 2.2e-08 |
| 6wnw_4 | 13 | 166 | RF00002 | 5_8S_rRNA | CL00112 | LSU | 1.7e-07 |
| 7aih_1 | 2,400 | 2,963 | RF02546 | LSU_trypano_mito | CL00112 | LSU | 8.6e-140 |
| 7aih_1 | 2,984 | 3,610 | RF02545 | SSU_trypano_mito | CL00111 | SSU | 3.5e-130 |
| 6ydp_AA | 1,176 | 2,737 | RF02541 | LSU_rRNA_bacteria | CL00112 | LSU | 2.5e-131 |
| 6ydp_AA | 1,173 | 2,738 | RF02540 | LSU_rRNA_archaea | CL00112 | LSU | 9.8e-92 |
| 6ydp_AA | 66 | 1,037 | RF00177 | SSU_rRNA_bacteria | CL00111 | SSU | 1e-80 |
| 6ydp_AA | 71 | 1,035 | RF01959 | SSU_rRNA_archaea | CL00111 | SSU | 2.2e-56 |
| 6ydp_AA | 71 | 1,032 | RF02542 | SSU_rRNA_microsporidia | CL00111 | SSU | 3.4e-46 |
| 6ydp_AA | 2,141 | 2,541 | RF02543 | LSU_rRNA_eukarya | CL00112 | LSU | 8e-33 |
| 6ydp_AA | 1,915 | 2,080 | RF02543 | LSU_rRNA_eukarya | CL00112 | LSU | 5.2e-12 |
| 6ydp_AA | 2,669 | 2,743 | RF00005 | tRNA | CL00001 | tRNA | 6.6e-10 |
| 6ydp_AA | 5,333 | 5,268 | RF00005 | tRNA | CL00001 | tRNA | 1.2e-09 |
| 6ydp_AA | 3,701 | 3,769 | RF00005 | tRNA | CL00001 | tRNA | 1.6e-09 |
| 6ydp_AA | 3,839 | 3,767 | RF00005 | tRNA | CL00001 | tRNA | 1.4e-08 |
| 6ydp_AA | 9,408 | 9,476 | RF00005 | tRNA | CL00001 | tRNA | 2.3e-07 |
| 6ydp_AA | 11,561 | 11,629 | RF00005 | tRNA | CL00001 | tRNA | 5.6e-07 |
| 6ydp_AA | 14,159 | 14,091 | RF00005 | tRNA | CL00001 | tRNA | 1.9e-06 |
| 6ydp_AA | 5,094 | 5,027 | RF00005 | tRNA | CL00001 | tRNA | 1e-05 |
| 6ydp_AA | 5,170 | 5,096 | RF00005 | tRNA | CL00001 | tRNA | 1.6e-05 |
| 6ydp_AA | 1 | 70 | RF00005 | tRNA | CL00001 | tRNA | 1.7e-05 |
| 6ydp_AA | 3,841 | 3,910 | RF00005 | tRNA | CL00001 | tRNA | 0.00028 |
| 6ydp_AA | 6,951 | 6,883 | RF00005 | tRNA | CL00001 | tRNA | 0.00041 |
| 7lyg_A | 32 | 102 | RF02340 | DENV_SLA | CL00129 | Flavivirus-5UTR | 2.8e-10 |
| 7lyg_A | 1 | 139 | RF00005 | tRNA | CL00001 | tRNA | 1e-09 |
| 6uz7_8 | 1 | 1,507 | RF01960 | SSU_rRNA_eukarya | CL00111 | SSU | 0.0 |
| 6uz7_8 | 1 | 1,507 | RF02542 | SSU_rRNA_microsporidia | CL00111 | SSU | 3.3e-238 |
| 6uz7_8 | 1 | 1,510 | RF01959 | SSU_rRNA_archaea | CL00111 | SSU | 1.6e-180 |
| 6uz7_8 | 1 | 1,512 | RF00177 | SSU_rRNA_bacteria | CL00111 | SSU | 1.2e-164 |
| 6uz7_8 | 2,140 | 2,825 | RF02543 | LSU_rRNA_eukarya | CL00112 | LSU | 4.6e-141 |
| 6uz7_8 | 1,724 | 2,825 | RF02541 | LSU_rRNA_bacteria | CL00112 | LSU | 8.6e-84 |
| 6uz7_8 | 1,975 | 2,825 | RF02540 | LSU_rRNA_archaea | CL00112 | LSU | 2.1e-71 |
| 6uz7_8 | 1,736 | 1,889 | RF00002 | 5_8S_rRNA | CL00112 | LSU | 9.9e-45 |
| 7lyf_A | 31 | 100 | RF02340 | DENV_SLA | CL00129 | Flavivirus-5UTR | 6.6e-16 |
| 7lyf_A | 31 | 139 | RF03546 | Flavivirus-5UTR | CL00129 | Flavivirus-5UTR | 8.8e-11 |
| 7lyf_A | 1 | 136 | RF00005 | tRNA | CL00001 | tRNA | 4.6e-10 |
| 8dfv_E | 1 | 57 | RF00027 | let-7 | CL00148 | let-7 | 3.6e-09 |
| 8dfv_E | 57 | 2 | RF00027 | let-7 | CL00148 | let-7 | 8.2e-09 |
| 8dfv_E | 56 | 1 | RF04289 | mir-3596 | CL00148 | let-7 | 1.1e-05 |
| 8dfv_E | 3 | 57 | RF04289 | mir-3596 | CL00148 | let-7 | 4.1e-05 |
| 8dfv_E | 58 | 1 | RF04292 | mir-379 | CL00149 | mir-154 | 0.00024 |

#### 2. Table of PDB RNA chains with homology to multiple Rfam families that do not share a common Rfam Clan

**Table S2:** PDB RNA chain cluster representatives (at 99% sequence similarity) with homology to multiple Rfam families that do not share a common Clan. We provide all Infernal `cmscan` hits at an E-value threshold of $10^{-3}$.

| RNA chain | Start | End | Rfam accession | Name | Clan accesion | Clan name | E-value |
|---|---|---|---|---|---|---|---|
| 7am2_1 | 1,904 | 2,470 | RF02546 | LSU_trypano_mito | CL00112 | LSU | 1.4e-138 |
| 7am2_1 | 2,491 | 3,117 | RF02545 | SSU_trypano_mito | CL00111 | SSU | 4e-129 |
| 1ser_T | 1 | 91 | RF00005 | tRNA | CL00001 | tRNA | 5.4e-15 |
| 1ser_T | 1 | 90 | RF01852 | tRNA-Sec | CL00001 | tRNA | 3.2e-08 |
| 1ser_T | 1 | 94 | RF02223 | sX4 | | | 4.6e-06 |
| 7nww_z | 1 | 85 | RF00005 | tRNA | CL00001 | tRNA | 2.4e-12 |
| 7nww_z | 1 | 88 | RF02223 | sX4 | | | 1.7e-05 |
| 1m5k_B | 1 | 69 | RF00173 | Hairpin | | | 8.6e-10 |
| 1m5k_B | 1 | 68 | RF04190 | Hairpin-meta1 | | | 1.4e-07 |
| 1m5k_B | 1 | 69 | RF04191 | Hairpin-meta2 | | | 3.3e-05 |
| 7nfx_1 | 1 | 299 | RF00017 | Metazoa_SRP | CL00003 | SRP | 1.7e-76 |
| 7nfx_1 | 1 | 299 | RF01855 | Plant_SRP | CL00003 | SRP | 5.1e-29 |
| 7nfx_1 | 17 | 276 | RF01856 | Protozoa_SRP | CL00003 | SRP | 1.3e-21 |
| 7nfx_1 | 1 | 297 | RF01857 | Archaea_SRP | CL00003 | SRP | 1.6e-09 |
| 7nfx_1 | 1 | 50 | RF04277 | mir-1268 | | | 2.4e-09 |
| 7nfx_1 | 14 | 289 | RF01570 | Dictyostelium_SRP | CL00003 | SRP | 1.1e-07 |
| 7obq_1 | 1 | 249 | RF00017 | Metazoa_SRP | CL00003 | SRP | 3.5e-58 |
| 7obq_1 | 1 | 249 | RF01855 | Plant_SRP | CL00003 | SRP | 1.3e-24 |
| 7obq_1 | 17 | 238 | RF01856 | Protozoa_SRP | CL00003 | SRP | 2.7e-21 |
| 7obq_1 | 1 | 50 | RF04277 | mir-1268 | | | 2e-09 |
| 7obq_1 | 60 | 238 | RF01857 | Archaea_SRP | CL00003 | SRP | 8.6e-08 |
| 7obq_1 | 67 | 231 | RF01570 | Dictyostelium_SRP | CL00003 | SRP | 6.2e-06 |
| 5aox_F | 2 | 86 | RF00017 | Metazoa_SRP | CL00003 | SRP | 9.8e-11 |
| 5aox_F | 39 | 1 | RF03639 | mir-619 | | | 2.4e-06 |
| 6mj0_B | 24 | 104 | RF00233 | Tymo_tRNA-like | | | 4.9e-16 |
| 6mj0_B | 1 | 23 | RF00390 | UPSK | | | 5.9e-07 |
| 6r6g_AF | 1 | 206 | RF00017 | Metazoa_SRP | CL00003 | SRP | 2.2e-41 |
| 6r6g_AF | 67 | 188 | RF01856 | Protozoa_SRP | CL00003 | SRP | 2.3e-14 |
| 6r6g_AF | 1 | 206 | RF01855 | Plant_SRP | CL00003 | SRP | 6.6e-13 |
| 6r6g_AF | 1 | 50 | RF04277 | mir-1268 | | | 1.7e-09 |
| 3jaj_4 | 1 | 206 | RF00017 | Metazoa_SRP | CL00003 | SRP | 2.7e-43 |
| 3jaj_4 | 1 | 206 | RF01855 | Plant_SRP | CL00003 | SRP | 2.6e-16 |
| 3jaj_4 | 17 | 206 | RF01856 | Protozoa_SRP | CL00003 | SRP | 9.7e-15 |
| 3jaj_4 | 1 | 50 | RF04277 | mir-1268 | | | 1.7e-09 |
| 3jaj_4 | 1 | 206 | RF01857 | Archaea_SRP | CL00003 | SRP | 0.00064 |
| 7sam_A | 39 | 167 | RF01084 | TLS-PK3 | | | 4.2e-28 |
| 7sam_A | 37 | 171 | RF01085 | TLS-PK4 | | | 8e-08 |
| 8fli_A | 77 | 226 | RF02012 | group-II-D1D4-7 | CL00102 | group-II-D1D4 | 7.4e-26 |
| 8fli_A | 807 | 884 | RF00029 | Intron_gpII | | | 3.2e-09 |
| 4v5z_BE | 4 | 51 | RF03852 | mir-610 | | | 7.4e-09 |
| 4v5z_BE | 51 | 4 | RF03852 | mir-610 | | | 7.4e-09 |
| 4v5z_BE | 1 | 54 | RF03934 | mir-m107-1 | | | 0.00038 |
| 4v5z_BE | 54 | 1 | RF03934 | mir-m107-1 | | | 0.00038 |
| 6chr_A | 548 | 621 | RF00029 | Intron_gpII | | | 3e-13 |
| 6chr_A | 42 | 199 | RF02012 | group-II-D1D4-7 | CL00102 | group-II-D1D4 | 1.9e-10 |
| 6chr_A | 25 | 177 | RF02003 | group-II-D1D4-4 | CL00102 | group-II-D1D4 | 9.7e-05 |
| 4ue5_A | 1 | 299 | RF00017 | Metazoa_SRP | CL00003 | SRP | 4e-76 |
| 4ue5_A | 1 | 299 | RF01855 | Plant_SRP | CL00003 | SRP | 1.9e-28 |
| 4ue5_A | 17 | 268 | RF01856 | Protozoa_SRP | CL00003 | SRP | 1.4e-21 |
| 4ue5_A | 1 | 297 | RF01857 | Archaea_SRP | CL00003 | SRP | 2e-09 |
| 4ue5_A | 1 | 50 | RF04277 | mir-1268 | | | 2.4e-09 |
| 4ue5_A | 14 | 283 | RF01570 | Dictyostelium_SRP | CL00003 | SRP | 3.8e-07 |
| 4ds6_A | 91 | 263 | RF02001 | group-II-D1D4-3 | CL00102 | group-II-D1D4 | 4.7e-31 |
| 4ds6_A | 261 | 330 | RF01998 | group-II-D1D4-1 | CL00102 | group-II-D1D4 | 1.8e-07 |
| 4ds6_A | 362 | 419 | RF00029 | Intron_gpII | | | 8.5e-05 |

| RNA chain | Start | End | Rfam accession | Name | Clan accesion | Clan name | E-value |
|---|---|---|---|---|---|---|---|
| 6hrm_1 | 1,507 | 4,403 | RF02541 | LSU_rRNA_bacteria | CL00112 | LSU | 0.0 |
| 6hrm_1 | 1,506 | 4,411 | RF02540 | LSU_rRNA_archaea | CL00112 | LSU | 0.0 |
| 6hrm_1 | 1 | 1,528 | RF00177 | SSU_rRNA_bacteria | CL00111 | SSU | 0.0 |
| 6hrm_1 | 1,666 | 4,414 | RF02543 | LSU_rRNA_eukarya | CL00112 | LSU | 0.0 |
| 6hrm_1 | 6 | 1,460 | RF01959 | SSU_rRNA_archaea | CL00111 | SSU | 3.5e-274 |
| 6hrm_1 | 6 | 1,457 | RF02542 | SSU_rRNA_microsporidia | CL00111 | SSU | 8.3e-199 |
| 6hrm_1 | 6 | 1,522 | RF01960 | SSU_rRNA_eukarya | CL00111 | SSU | 8.4e-181 |
| 6hrm_1 | 1,519 | 1,672 | RF00002 | 5_8S_rRNA | CL00112 | LSU | 2.5e-07 |
| 6hrm_1 | 4,372 | 4,445 | RF00177 | SSU_rRNA_bacteria | CL00111 | SSU | 0.00072 |
| 8dvs_B | 1 | 64 | RF04070 | MIR6440 | | | 2.1e-05 |
| 8dvs_B | 64 | 1 | RF04070 | MIR6440 | | | 3.1e-05 |
| 8dvs_B | 3 | 62 | RF03819 | mir-Ro6-3 | | | 4.2e-05 |
| 8dvs_B | 62 | 3 | RF03819 | mir-Ro6-3 | | | 4.3e-05 |
| 8dvs_B | 64 | 1 | RF00729 | mir-278 | | | 5.1e-05 |
| 8dvs_B | 5 | 60 | RF03934 | mir-m107-1 | | | 8e-05 |
| 8dvs_B | 64 | 1 | RF03771 | mir-341 | | | 9.5e-05 |
| 8dvs_B | 60 | 5 | RF03934 | mir-m107-1 | | | 0.00011 |
| 8dvs_B | 1 | 64 | RF03771 | mir-341 | | | 0.00012 |
| 8dvs_B | 2 | 63 | RF04026 | mir-8499 | | | 0.00023 |
| 8dvs_B | 63 | 2 | RF04026 | mir-8499 | | | 0.00024 |
| 8dvs_B | 62 | 3 | RF01688 | Actino-pnp | | | 0.00037 |
| 8gzq_S | 2 | 70 | RF02340 | DENV_SLA | CL00129 | Flavivirus-5UTR | 9.2e-18 |
| 8gzq_S | 74 | 168 | RF00185 | Flavi_CRE | | | 1.8e-09 |
| 8gzq_S | 2 | 113 | RF03546 | Flavivirus-5UTR | CL00129 | Flavivirus-5UTR | 4.6e-06 |
| 7aor_2 | 2,020 | 2,579 | RF02546 | LSU_trypano_mito | CL00112 | LSU | 9.7e-138 |
| 7aor_2 | 2,589 | 3,210 | RF02545 | SSU_trypano_mito | CL00111 | SSU | 4.5e-120 |
| 4ujd_BC | 2 | 354 | RF00061 | IRES_HCV | CL00017 | IRES1 | 5.4e-138 |
| 4ujd_BC | 128 | 321 | RF00209 | IRES_Pesti | CL00017 | IRES1 | 2.6e-10 |
| 4ujd_BC | 387 | 485 | RF00620 | HCV_ARF_SL | | | 2.6e-06 |
| 8t2s_B | 101 | 278 | RF02001 | group-II-D1D4-3 | CL00102 | group-II-D1D4 | 1.5e-28 |
| 8t2s_B | 573 | 652 | RF00029 | Intron_gpII | | | 4.1e-08 |
| 8t2s_B | 315 | 393 | RF01998 | group-II-D1D4-1 | CL00102 | group-II-D1D4 | 5e-06 |
| 6wnw_4 | 1 | 3,031 | RF02541 | LSU_rRNA_bacteria | CL00112 | LSU | 0.0 |
| 6wnw_4 | 1 | 3,031 | RF02540 | LSU_rRNA_archaea | CL00112 | LSU | 0.0 |
| 6wnw_4 | 160 | 3,021 | RF02543 | LSU_rRNA_eukarya | CL00112 | LSU | 0.0 |
| 6wnw_4 | 1,069 | 1,185 | RF00001 | 5S_rRNA | CL00113 | 5S_rRNA | 2.2e-08 |
| 6wnw_4 | 13 | 166 | RF00002 | 5_8S_rRNA | CL00112 | LSU | 1.7e-07 |
| 5lzf_x | 13 | 48 | RF01988 | SECIS_2 | | | 1.2e-06 |
| 5lzf_x | 15 | 48 | RF01989 | SECIS_3 | | | 6.6e-05 |
| 8s95_C | 32 | 122 | RF00386 | Entero_5_CRE | | | 8.4e-26 |
| 8s95_C | 2 | 154 | RF00005 | tRNA | CL00001 | tRNA | 1.3e-09 |
| 8fti_B | 3 | 67 | RF04036 | mir-2076 | | | 2.4e-08 |
| 8fti_B | 64 | 2 | RF00827 | mir-77 | | | 9.6e-06 |
| 8fti_B | 59 | 10 | RF03658 | mir-2300 | | | 1.7e-05 |
| 8fti_B | 10 | 59 | RF03658 | mir-2300 | | | 1.7e-05 |
| 8fti_B | 67 | 2 | RF04294 | mir-3578 | CL00149 | mir-154 | 3.6e-05 |
| 8fti_B | 7 | 62 | RF03257 | mir-4427 | | | 4.8e-05 |
| 8fti_B | 66 | 3 | RF03404 | mir-1298 | | | 8.4e-05 |
| 8fti_B | 3 | 66 | RF03404 | mir-1298 | | | 0.00011 |
| 8fti_B | 1 | 69 | RF04295 | mir-329 | CL00149 | mir-154 | 0.00012 |
| 8fti_B | 11 | 58 | RF03938 | mir-4662 | | | 0.00014 |
| 8fti_B | 6 | 65 | RF03601 | MIR7993 | | | 0.00019 |
| 8fti_B | 1 | 72 | RF03743 | MIR7696 | | | 0.00019 |
| 8fti_B | 70 | 2 | RF00928 | mir-590 | | | 0.00029 |
| 8fti_B | 76 | 1 | RF03743 | MIR7696 | | | 0.0003 |
| 8fti_B | 62 | 7 | RF03257 | mir-4427 | | | 0.00089 |
| 1e8s_C | 21 | 88 | RF00017 | Metazoa_SRP | CL00003 | SRP | 1.7e-10 |
| 1e8s_C | 19 | 70 | RF04277 | mir-1268 | | | 2.6e-10 |
| 1e8s_C | 21 | 88 | RF01855 | Plant_SRP | CL00003 | SRP | 0.00012 |
| 1grz_B | 1 | 237 | RF00028 | Intron_gpI | | | 7.4e-30 |
| 1grz_B | 125 | 246 | RF03000 | LOOT | | | 0.00048 |
| 6q97_7 | 1 | 74 | RF00005 | tRNA | CL00001 | tRNA | 2.8e-15 |
| 6q97_7 | 77 | 39 | RF02194 | HPnc0260 | | | 2.4e-06 |
| 7aih_1 | 2,400 | 2,963 | RF02546 | LSU_trypano_mito | CL00112 | LSU | 8.6e-140 |
| 7aih_1 | 2,984 | 3,610 | RF02545 | SSU_trypano_mito | CL00111 | SSU | 3.5e-130 |
| 7lyf_A | 31 | 100 | RF02340 | DENV_SLA | CL00129 | Flavivirus-5UTR | 6.6e-16 |
| 7lyf_A | 31 | 139 | RF03546 | Flavivirus-5UTR | CL00129 | Flavivirus-5UTR | 8.8e-11 |
| 7lyf_A | 1 | 136 | RF00005 | tRNA | CL00001 | tRNA | 4.6e-10 |

| RNA chain | Start | End | Rfam accession | Name | Clan accesion | Clan name | E-value |
|---|---|---|---|---|---|---|---|
| 8sp9_C | 30 | 119 | RF00386 | Entero_5_CRE | | | 4.5e-28 |
| 8sp9_C | 1 | 150 | RF00005 | tRNA | CL00001 | tRNA | 1.1e-07 |
| 4v5z_BF | 8 | 113 | RF03296 | MIR8788 | | | 3.8e-10 |
| 4v5z_BF | 113 | 8 | RF03296 | MIR8788 | | | 3.8e-10 |
| 4v5z_BF | 2 | 119 | RF04104 | MIR918 | | | 6e-10 |
| 4v5z_BF | 119 | 2 | RF04104 | MIR918 | | | 6e-10 |
| 4v5z_BF | 18 | 104 | RF04248 | MIR7486 | | | 5.4e-09 |
| 4v5z_BF | 103 | 17 | RF04248 | MIR7486 | | | 5.4e-09 |
| 4v5z_BF | 32 | 89 | RF04076 | mir-H11 | | | 2.8e-07 |
| 4v5z_BF | 89 | 32 | RF04076 | mir-H11 | | | 2.8e-07 |
| 4v5z_BF | 20 | 101 | RF03703 | mir-3622 | | | 4.8e-07 |
| 4v5z_BF | 101 | 20 | RF03703 | mir-3622 | | | 4.8e-07 |
| 4v5z_BF | 10 | 111 | RF03976 | MIR898 | | | 8.5e-05 |
| 4v5z_BF | 111 | 10 | RF03976 | MIR898 | | | 8.5e-05 |
| 4v5z_BF | 21 | 99 | RF04162 | MIR9662 | | | 0.00038 |
| 4v5z_BF | 100 | 22 | RF04162 | MIR9662 | | | 0.00038 |
| 6ydp_AA | 1,176 | 2,737 | RF02541 | LSU_rRNA_bacteria | CL00112 | LSU | 2.5e-131 |
| 6ydp_AA | 1,173 | 2,738 | RF02540 | LSU_rRNA_archaea | CL00112 | LSU | 9.8e-92 |
| 6ydp_AA | 66 | 1,037 | RF00177 | SSU_rRNA_bacteria | CL00111 | SSU | 1e-80 |
| 6ydp_AA | 71 | 1,035 | RF01959 | SSU_rRNA_archaea | CL00111 | SSU | 2.2e-56 |
| 6ydp_AA | 71 | 1,032 | RF02542 | SSU_rRNA_microsporidia | CL00111 | SSU | 3.4e-46 |
| 6ydp_AA | 2,141 | 2,541 | RF02543 | LSU_rRNA_eukarya | CL00112 | LSU | 8e-33 |
| 6ydp_AA | 1,915 | 2,080 | RF02543 | LSU_rRNA_eukarya | CL00112 | LSU | 5.2e-12 |
| 6ydp_AA | 2,669 | 2,743 | RF00005 | tRNA | CL00001 | tRNA | 6.6e-10 |
| 6ydp_AA | 5,333 | 5,268 | RF00005 | tRNA | CL00001 | tRNA | 1.2e-09 |
| 6ydp_AA | 3,701 | 3,769 | RF00005 | tRNA | CL00001 | tRNA | 1.6e-09 |
| 6ydp_AA | 3,839 | 3,767 | RF00005 | tRNA | CL00001 | tRNA | 1.4e-08 |
| 6ydp_AA | 9,408 | 9,476 | RF00005 | tRNA | CL00001 | tRNA | 2.3e-07 |
| 6ydp_AA | 11,561 | 11,629 | RF00005 | tRNA | CL00001 | tRNA | 5.6e-07 |
| 6ydp_AA | 14,159 | 14,091 | RF00005 | tRNA | CL00001 | tRNA | 1.9e-06 |
| 6ydp_AA | 5,094 | 5,027 | RF00005 | tRNA | CL00001 | tRNA | 1e-05 |
| 6ydp_AA | 5,170 | 5,096 | RF00005 | tRNA | CL00001 | tRNA | 1.6e-05 |
| 6ydp_AA | 1 | 70 | RF00005 | tRNA | CL00001 | tRNA | 1.7e-05 |
| 6ydp_AA | 3,841 | 3,910 | RF00005 | tRNA | CL00001 | tRNA | 0.00028 |
| 6ydp_AA | 6,951 | 6,883 | RF00005 | tRNA | CL00001 | tRNA | 0.00041 |
| 6ydp_AA | 16,450 | 16,378 | RF04217 | mir-297 | | | 0.0005 |
| 7lyg_A | 32 | 102 | RF02340 | DENV_SLA | CL00129 | Flavivirus-5UTR | 2.8e-10 |
| 7lyg_A | 1 | 139 | RF00005 | tRNA | CL00001 | tRNA | 1e-09 |
| 1e8o_E | 3 | 48 | RF00017 | Metazoa_SRP | CL00003 | SRP | 4.6e-09 |
| 1e8o_E | 1 | 50 | RF04277 | mir-1268 | | | 2.6e-08 |
| 1e8o_E | 3 | 48 | RF01855 | Plant_SRP | CL00003 | SRP | 0.00041 |
| 2zue_B | 1 | 75 | RF00005 | tRNA | CL00001 | tRNA | 2.7e-15 |
| 2zue_B | 1 | 74 | RF01852 | tRNA-Sec | CL00001 | tRNA | 8.3e-05 |
| 2zue_B | 19 | 74 | RF01684 | mascRNA-menRNA | | | 0.00021 |
| 8uw3_B | 55 | 1 | RF01940 | hvt-mir-H | | | 0.00011 |
| 8uw3_B | 59 | 1 | RF03516 | MIR5229 | | | 0.00014 |
| 8uw3_B | 54 | 1 | RF03654 | MIR4239 | | | 0.00032 |
| 8uw3_B | 61 | 1 | RF03872 | MIR6426 | | | 0.00081 |
| 8uw3_B | 72 | 1 | RF02481 | GlsR18 | | | 0.00095 |
| 3add_D | 1 | 88 | RF01852 | tRNA-Sec | CL00001 | tRNA | 7.3e-15 |
| 3add_D | 2 | 88 | RF00005 | tRNA | CL00001 | tRNA | 0.00018 |
| 3add_D | 20 | 87 | RF01684 | mascRNA-menRNA | | | 0.00032 |
| 5f9r_A | 38 | 116 | RF02348 | tracrRNA | | | 1e-12 |
| 5f9r_A | 50 | 21 | RF01335 | CRISPR-DR22 | | | 4.8e-05 |
| 5f9r_A | 37 | 66 | RF01335 | CRISPR-DR22 | | | 0.00048 |
| 6uz7_8 | 1 | 1,507 | RF01960 | SSU_rRNA_eukarya | CL00111 | SSU | 0.0 |
| 6uz7_8 | 1 | 1,507 | RF02542 | SSU_rRNA_microsporidia | CL00111 | SSU | 3.3e-238 |
| 6uz7_8 | 1 | 1,510 | RF01959 | SSU_rRNA_archaea | CL00111 | SSU | 1.6e-180 |
| 6uz7_8 | 1 | 1,512 | RF00177 | SSU_rRNA_bacteria | CL00111 | SSU | 1.2e-164 |
| 6uz7_8 | 2,140 | 2,825 | RF02543 | LSU_rRNA_eukarya | CL00112 | LSU | 4.6e-141 |
| 6uz7_8 | 1,724 | 2,825 | RF02541 | LSU_rRNA_bacteria | CL00112 | LSU | 8.6e-84 |
| 6uz7_8 | 1,975 | 2,825 | RF02540 | LSU_rRNA_archaea | CL00112 | LSU | 2.1e-71 |
| 6uz7_8 | 1,736 | 1,889 | RF00002 | 5_8S_rRNA | CL00112 | LSU | 9.9e-45 |
| 4v5z_BI | 5 | 68 | RF03976 | MIR898 | | | 1.1e-06 |
| 4v5z_BI | 68 | 5 | RF03976 | MIR898 | | | 1.1e-06 |
| 4v5z_BI | 9 | 65 | RF04248 | MIR7486 | | | 9.2e-06 |
| 4v5z_BI | 64 | 8 | RF04248 | MIR7486 | | | 9.2e-06 |
| 4v5z_BI | 1 | 72 | RF03819 | mir-Ro6-3 | | | 1.2e-05 |
| 4v5z_BI | 72 | 1 | RF03819 | mir-Ro6-3 | | | 1.2e-05 |
| 6wkr_H | 65 | 256 | RF04159 | MIR1437 | | | 9.2e-24 |
| 6wkr_H | 256 | 65 | RF04159 | MIR1437 | | | 2e-23 |
| 6wkr_H | 205 | 115 | RF04162 | MIR9662 | | | 1e-07 |
| 6wkr_H | 116 | 206 | RF04162 | MIR9662 | | | 2.4e-07 |
| 6wkr_H | 200 | 121 | RF03691 | mir-2765 | | | 4.5e-05 |
